# The arrival of the Near Eastern ancestry in Central Italy predates the onset of the Roman Empire

**DOI:** 10.1101/2024.10.07.617003

**Authors:** Francesco Ravasini, Cecilia Conati Barbaro, Christiana Lyn Scheib, Kristiina Tambets, Mait Metspalu, Fulvio Cruciani, Beniamino Trombetta, Eugenia D’Atanasio

## Abstract

Italian genetic history was profoundly shaped by Romans. While the Iron Age was comparable to contemporary European regions, the gene pool of Central Italy underwent significant influence from Near Eastern ancestry during the Imperial age. To explain this shift, it has been proposed that during this period people from Eastern Mediterranean regions of the Empire migrated towards its political center. In this study, by analyzing a new individual (1.25x) and published Republican samples, we propose a novel perspective for the presence of Near Eastern ancestry in the Imperial gene pool. In our scenario, the spread of this genetic ancestry took place during the late Republican period, predating the onset of the Empire by ∼200 years. The diffusion of this ancestry may have occurred due to early East-to-West movements, since Eastern Mediterranean regions were already under Roman political influence during the Republic, or even as a result of migration from Southern Italy where Greeks and Phoenicians settled.

## Introduction

The Italian peninsula, due to its central position, has always been a melting pot for the people living across the whole Mediterranean sea (Abulafia 2011; Broodbank 2013). The study of ancient DNA (aDNA) over the last 20 years greatly enhanced our comprehension of the movements of individuals within Italy and their impact on the local gene pool (Aneli et al. 2021). Overall, all these migrations resulted in a continuous shift in the genetic landscape of the peninsula, starting from the Paleolithic (Posth et al. 2023) to the modern times (Antonio et al. 2019; Raveane et al. 2019; Posth et al. 2021). Although we are still lacking the genetic characterization of several regions in different periods, necessary to portray a comprehensive picture, the evolution of the Italian gene pool seems to be characterized by periods of relative genetic diversity, both at the regional and the individual level (Antonio et al. 2019; Aneli et al. 2021; Posth et al. 2021; Saupe et al. 2021; Aneli et al. 2022; Ravasini et al. 2024).

A pivotal role in the homogenization of Italy, both from a cultural and genetic perspective, has been played by Romans. While the people that inhabited the Italian peninsula during the previous Iron Age (IA) were characterized by a great cultural diversity (Pallottino 1991; Bietti Sestieri 2010) and a relatively low genetic diversity (Antonio et al. 2019; Posth et al. 2021; Aneli et al. 2022; Ravasini et al. 2024), with the conquest of the peninsula, the Romans imposed their culture, language, institutions and social aspects, eventually influencing even the evolution of the gene pool (Antonio et al. 2019; Posth et al. 2021; Scorrano et al. 2022; Coia et al. 2023; Moots et al. 2023; Antonio et al. 2024; Ravasini et al. 2024).

The spread of the Roman domination began during the Republican period (509– 27 BCE), a fundamental stage of the history of Rome in which the small city-state became a hegemonic power, not only in Italy, but across the whole Mediterranean Sea. Among the many people and ethnicities subjected during their conquest of the peninsula, the Romans asserted their power over *Magna Graecia*, the Greek colonies established from the 8^th^ century BCE in Southern Italy, which reached elevated levels of political, cultural and societal development, even competing with their motherland (Brizzi 1997; van Dommelen 2012; De Angelis 2016; Hodos 2020). The political integration of *Magna Graecia* greatly influenced Roman society, eventually resulting in major cultural transformations. In the next centuries, the territories conquered and embedded in the Roman state steadily increased, coming into close contact with different people which, like the Southern Italy Greeks, contributed to the enrichment of the social substrate of Roman society. At the onset of the Empire (27 BCE), Rome was the most powerful and extensive European political entity, a cosmopolitan state encompassing the Mediterranean area.

Besides all the influence over politics and culture, the Roman Empire (27 BCE – 476 CE in Italy) had a great impact also on the genomic landscape of the conquered territories, especially in Italy. While the Italian Iron Age populations prior to the expansion of the Roman Republic were more similar to modern Northern Italians and Central Europeans (aside from small regional differences), in Imperial time a shift towards Near Eastern ancestries can be observed (Antonio et al. 2019; Posth et al. 2021; Aneli et al. 2022; Scorrano et al. 2022; Coia et al. 2023; Ravasini et al. 2024). This great Near Eastern genetic influence has been mostly explained as the result of massive migration, during Imperial time, from the Eastern Mediterranean regions into Rome (Antonio et al. 2019; Lazaridis et al. 2022) and other regions of the peninsula (Posth et al. 2021). In this context, it has been proposed that the high population density of Eastern Imperial provinces and/or the attractiveness of a power center like Rome may have been the cause of this process (Scheidel et al. 2007; Antonio et al. 2019).

Nevertheless, some considerations about the available historical and genetic data may point to an earlier arrival of Near Eastern ancestry in the italian peninsula which seems not to be a direct consequence of the onset of the Roman Empire. First, Rome annexed some of the rich and densely populated Near Eastern territories decades, if not centuries, before the onset of the Empire (Macedonia and Greece between 168 and 146 BCE; Western part of Anatolia in 133 BCE and soon afterwards Cilicia, the Southern part of Anatolia, from 100 BCE; and Syria in 64 BCE (Piganiol 1927; Rinaldi Tufi et al. 1971; Brizzi 1997)) suggesting that migrations from those regions might have started much earlier. Second, the individuals genetically analyzed so far dated to the very early years of the Empire have already a considerable amount of Near Eastern ancestry. For example, the first genome ever retrieved from the archaeological site of Pompeii (Scorrano et al. 2022) (which, due to the eruption that destroyed the city, is precisely dated to the 79 CE) clusters with other Imperial individuals of the subsequent centuries and it has a high proportion of Iran Neolithic genetic component (usually a good proxy for Near Eastern ancestry in Europe) (Scorrano et al. 2022). Although not impossible, it seems unlikely that in just a few generations since the onset of the Empire, migrations bringing the Near Eastern ancestry already had an impact on the Italian gene pool, regardless of the geographic origin of the sample. On these bases, it is important to note that the Near Eastern genetic component may have arrived in Central-Northern Italy as a result of internal migrations after the conquest of genetically understudied Southern Italy and Sicily IA. Indeed, we are still lacking an extensive genomic characterization of *Magna Graecia* and Punic Sicily individuals, who probably had a Near Eastern ancestry since they arrived in Italy from the Eastern Mediterranean. Finally, among the few analyzed individuals dated to the latest period of the Roman Republic (the last two centuries of the 1^st^ millennium BCE) there are several ones interpreted as “genetic outliers” with a Levantine or Eastern Mediterranean putative origin (Antonio et al. 2019; Posth et al. 2021; Moots et al. 2023). These individuals may be the direct representatives of the ongoing arrival of Near Eastern ancestry which later characterized the genomic landscape of the Imperial period.

To better shed light on this fundamental shift of Italian gene pool towards the Near East, we performed a medium coverage sequencing (1.25x) of a late Republican individual from the site of Villa Falgari, located in Tarquinia, Viterbo (only 70 km away from Rome, Figure 1), who may be ascribed as a Levantine “genetic outlier”. By analyzing our new sample in the context of coeval individuals with similar genetic makeup, we show that the gene flow carrying the Near Eastern ancestry was probably already occurring during the late Republican phase, possibly indicating these “genetic outliers” as the individuals upon which the Imperial gene pool was founded.

**Figure 1.**
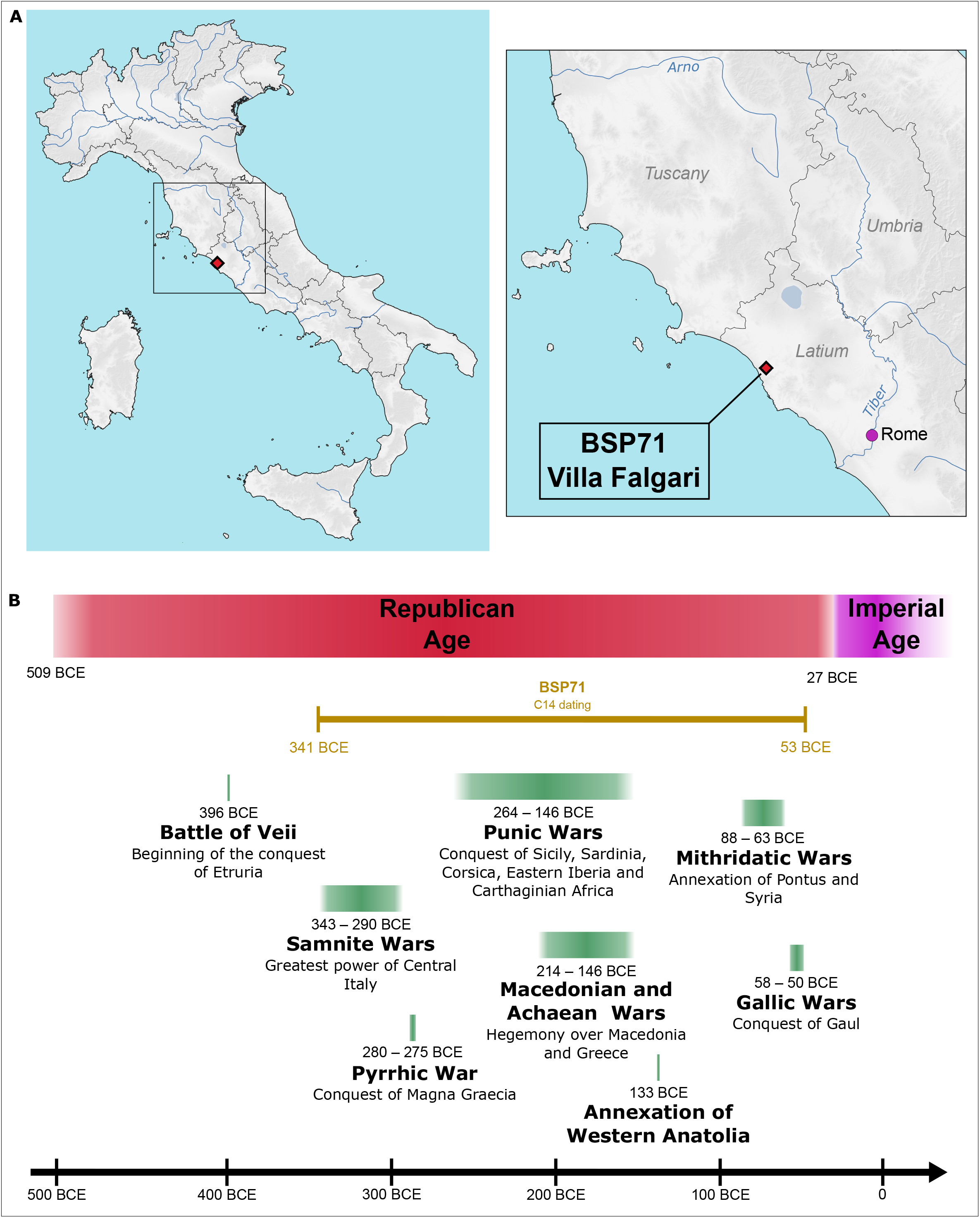
A) Geographic location of Villa Falgari site. B) Timeline of the Roman Republic and the first years of the Empire. Some of the main wars and annexations of the Republic are indicated. Two sigma calibrated radiocarbon dating of the sample BSP71 from Villa Falgari is portrayed in ochre.

## Results

We extracted DNA from an individual (named BSP71) found in the site of Villa Falgari, Tarquinia (Figure 1A; Supplementary Table 1) (Barich et al. 1968). Radiocarbon dating places this individual in the Republican period (341-53 2σ calBCE, 2126 ± 24 BP), more likely in the very last phase of the Republic, after the beginning of Roman expansionism in the Eastern Mediterranean (Figure 1; Supplementary table 2). Shotgun sequencing was performed resulting in a depth of coverage of 1.25x. The sequences were confirmed to be mostly ancient with negligible amount of contamination, and a male genetic sex was assigned for this individual (Supplementary table 1, Supplementary Methods). Genotype data were merged with ancient and modern individuals included in the Allen Ancient DNA Resource (AADR, v54.1 (Dataverse 7.0) Nov 16 2022 https://doi.org/10.7910/DVN/FFIDCW; Mallick et al. 2024) and in Reitsema et al. (2022) (Supplementary table 3).

PCA was performed to contextualize the Villa Falgari site in the genetic variability of Western Eurasia and North Africa. BSP71 results to be not included in the main cluster of IA/Republican individuals, being strongly shifted towards Near Eastern ancient and modern individuals and overlapping with the genetic variability of the subsequent Imperial period (Figure 2A, Supplementary figure 1). Interestingly, other IA/Republican individuals reported in the literature show this kind of genetic shift (Figure 2A, Supplementary figure 1) (Antonio et al. 2019; Moots et al. 2023). In particular, two individuals from Tarquinia (only a few kilometers far from Villa Falgari and dated to a similar late Republican phase, in the first two centuries BCE) are placed in the most Eastern extreme of the distribution, resulting very similar to IA Near Eastern groups like Phoenicians (Moots et al. 2023) (Figure 2A, Supplementary figure 1). Admixture analysis (Alexander et al. 2009) reflects these findings, showing a high proportion of Iran_N/CHG (i.e., Iran Neolithic/Caucasus Hunter-Gatherer) genetic component in aforementioned individuals and in Villa Falgari, while it has negligible proportion in the main IA/Republic cluster (Figure 2B, Supplementary figure 2). Very similar genetic make-up can be observed during the Imperial time, suggesting some kind of genetic connections between this period and the IA/Republican samples with Near Eastern ancestries (hereafter indicated as IA/Republic oNE) (Figure 2B). Similarly, D-statistics confirms this possible genetic link. When it is in the form D(IA/Republican oNE or Imperial groups, IA/Republic main cluster; Iran_N, ONG.SG) it shows that the first groups always share more derived alleles with Iran_N compared to IA/Republic main cluster. On the other hand, when Imperial and IA/Republic oNE groups are compared (D(IA/Republican oNE, Imperial groups; Iran_N, ONG.SG)), no significant difference can be observed, indicating a similar influence of this ancestry in these groups (Supplementary table 4). These results suggest that IA/Republican oNE and Imperial groups form a homogeneous genetic cluster in their affinity to Iran_N when they are compared to the IA/Republican main cluster.

**Figure 2.**
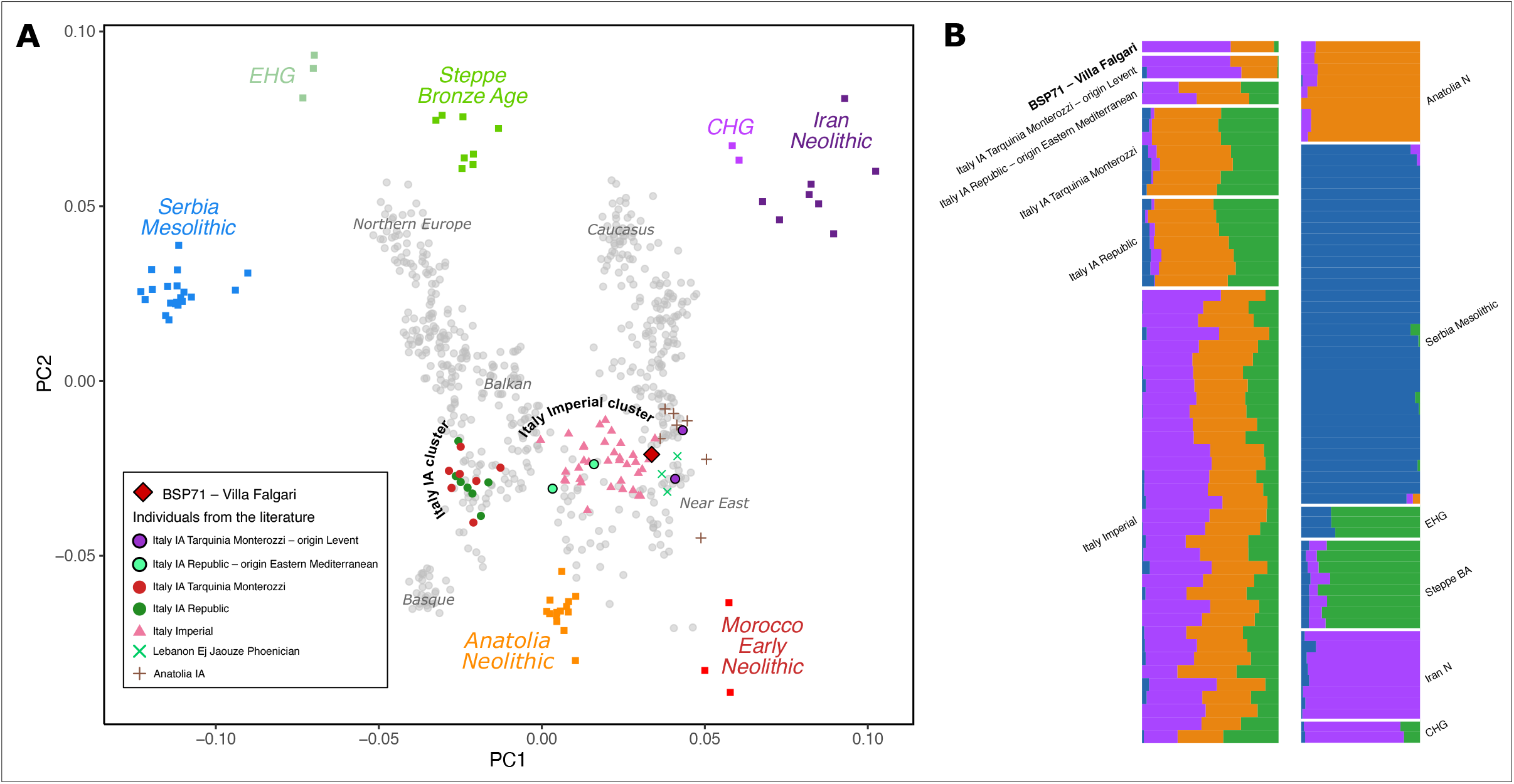
A) PCA with the newly reported individual and relevant ancient (colored) and modern (gray) samples from the literature. The main Italian Iron Age and Imperial clusters are indicated together with the Iron Age individuals with Near Eastern ancestry. B) Unsupervised admixture analysis (K=4), on the left Italian Iron Age and Imperial individuals are represented; on the right, populations representative of the main European genetic ancestries.

Interestingly, also the Y chromosome analysis comparing ancient individuals with modern ones shows clues of a Near Eastern ancestry in BSP71 and the other four IA/Republic oNE (R437, R10337, R10341, and R850) individuals (Supplementary figure 3 and supplementary table 5) (Karmin et al. 2015; Mallick et al. 2016; Poznik et al. 2016; Martiniano et al. 2022). BSP71 belongs to the R1-M269/Z2105 haplogroup and clusters with several Eastern individuals, mostly from late Bronze Age/Iron Age Armenia. The R437 individual belongs to a sister clade (R1-M269/P312), now frequent in Western Europe but also observed in Bronze Age Croatians. R10337 belongs to the R2-M479 clade, observed in present-day Western Asian groups and in ancient Middle Eastern people. Finally, R10341 and R850 belong to the J1-M267/P58 and T-M70 haplogroups respectively. These lineages are now observed mostly in Middle Eastern populations, but they were also present among ancient Eastern Mediterranean individuals, mostly from Greece and Anatolia.

**Figure 3.**
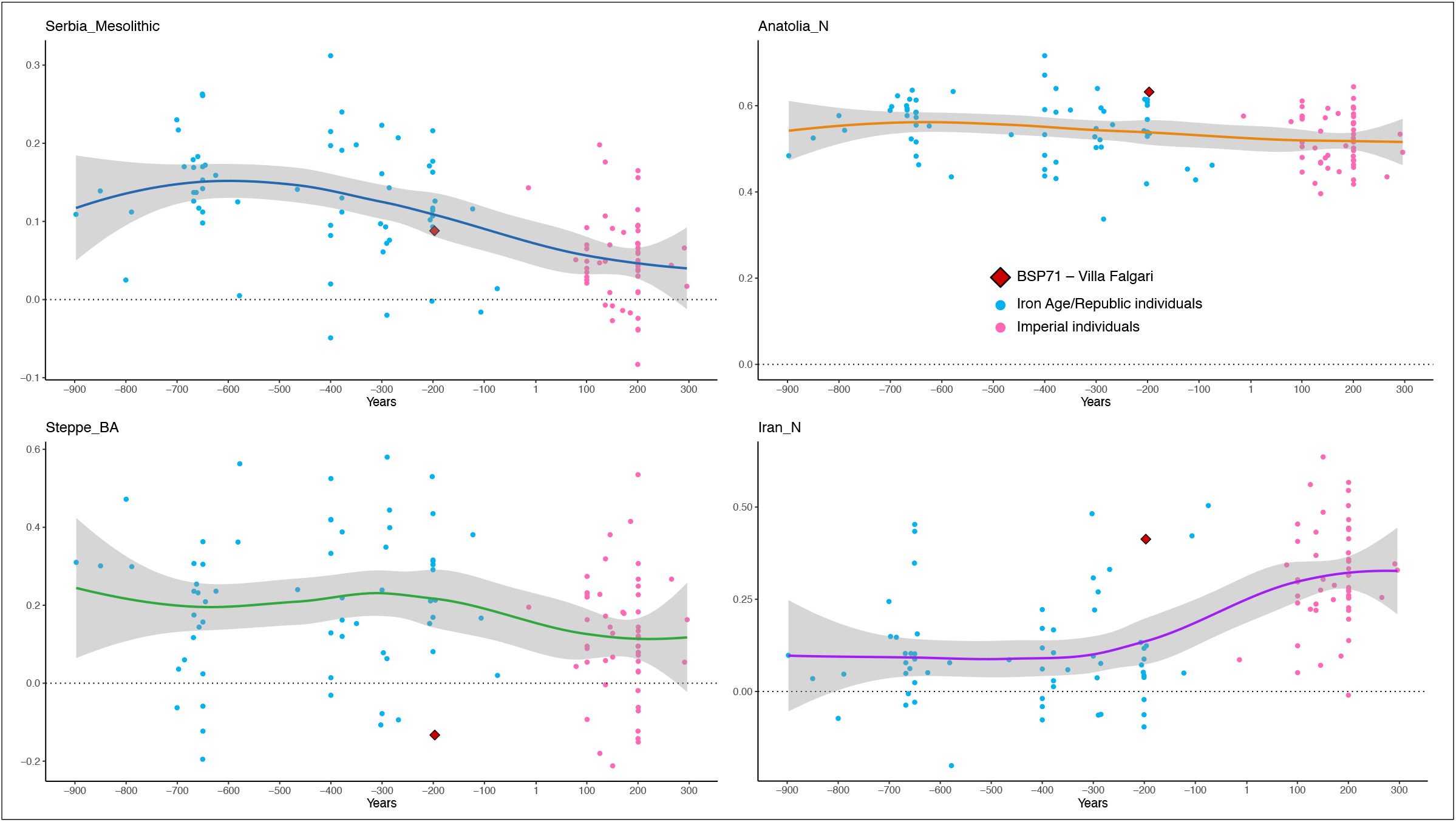
qpAdm analysis performed on Central Italian individuals from 900 BCE to 300 CE using the four sources: Serbia Mesolithic, Anatolia Neolithic, Steppe Bronze Age and Iran Neolithic. A sharp increase of the Iran Neolithic source can be observed starting from ∼200 BCE.

We explored more in detail these results by performing different tests of qpAdm modeling (Harney et al. 2021). Both the IA/Republic oNE and the Imperial groups can be modeled with a two source model including a main cluster IA/Republican (e.g., Italy_IA_Republic.SG) and a Near Eastern group (e.g., Turkey_IA) (Supplementary table 6). This finding confirms the strong Near Eastern influence in these groups, likely indicating that their gene pool emerged thanks to the gene flow from Near Eastern regions into Central Italy. More interestingly, the Imperial groups can be modeled as a mixture between main cluster IA/Republican groups (e.g., Italy_IA_Republic.SG) and the IA/Republic oNE ones (Supplementary table 7), indicating that the Imperial and subsequent gene pools may have arisen by admixture events between individuals similar to the IA/Republic oNE arrived during the Republican phase and the previous local gene pool. This would eventually indicate that the Near Eastern genetic influence in Central Italy started to spread during the Republic rather than at the onset of the Empire.

We further investigated this possibility by performing qpAdm on single individuals from Central Italy dated between 900 BCE to 300 CE (therefore encompassing the early IA, the Republican and first Imperial period) using 4 source populations which contributed to the peopling of Europe (Serbia_Mesolithic, Anatolia_N, Steppe_BA and Iran_N; Figure 3 and Supplementary table 8). While the Anatolia_N component remains stable over the centuries, the Iran_N one sharply increases starting from 200 BCE at the expense of the Steppe_BA and the Serbia_Mesolithic components (Figure 3). Despite the small number of samples belonging to the first two centuries BCE, this result may suggest that it is possible to frame the beginning of the spread of Near Eastern ancestry in Central Italy from the late Republican phase rather than the Imperial period. Nevertheless, it is very likely that this process lasted throughout the subsequent periods, at least to the early stages of the Empire. Indeed, it is possible to model Imperial groups as a mixture between several IA/Republic oNE groups (e.g., Villa Falgari) and Near Eastern populations, suggesting that an additional genetic influence from those regions arrived in Italy during the Imperial period (Supplementary Table 9).

A possible direct estimation of this admixture time was performed with DATES (Chintalapati et al. 2022), using the sources Italy_IA_Republic.SG (as a proxy of the original Central Italian ancestry) and Armenia_LBA_EIA (as a proxy of the Near Eastern ancestry). For Villa Falgari the estimated date resulted to be 5.043 ± 3.302 generations, corresponding to 141.204 ± 92.456 years before the age of the sample (considering 28 years for generation), while for the Italy_Imperial.SG group resulted to be 16.028 ± 8.141 generations, therefore 448.784 ± 227.948 years before the age of the sample. Considering the average dating for these groups, the admixture dates are in a similar range between the 4^th^ and the 2^nd^ century BCE, compatible with the dates obtained with different methods.

Finally, we performed an outgroup f3-statistic in the form f3(BSP71, *Test*; ONG.SG) where *Tests* are Eurasian groups comprising a 1,000 year range within the midpoint date of Villa Falgari individual (therefore including Imperial individuals) or modern populations (Supplementary figure 4 and supplementary table 10). This analysis shows that for ancient populations, the area with greater genetic similarity to BSP71 is the Central-Eastern Mediterranean, while with modern populations this area shifts towards Central-Southern Europe, suggesting that individuals like BSP71 may represent the first step of the modern gene pool of the area.

## Discussion and Conclusions

In this study we analyzed a late Roman Republican individual from the site of Villa Falgari and, reanalyzing published individuals from the same area with similar genetic make-up, we provided new insights into the diffusion of Near Eastern ancestry in Central Italy. Given the presence of a high proportion of Near Eastern ancestry in late Republican individuals, here we propose that the observed shift in Central Italian gene pool towards Eastern genetic components was not a consequence of the establishment of the Roman Empire, but an ongoing process started probably 200 years before the onset of the Empire. Movements of people from Eastern regions may have already occurred after the end of the Punic Wars, when Rome became the hegemonic power of the Mediterranean Sea and they probably increased when Macedonia, Greece and Anatolia were annexed to the Republic in the first two centuries BCE. Moreover, it is even plausible that the Near Eastern genetic influence arrived earlier in Central Italy in an indirect way, as a consequence of migration within Italy. Indeed, the conquest of Southern Italy, divided between *Magna Graecia* and Phoenicians, may have given rise to the spread of individuals who already carried Near Eastern ancestry.

Although at the moment few IA samples from Southern Italy have been analyzed, some hints suggest that their gene pool was influenced by the Near East. Middle and late Bronze Age individuals from Sicily already show the presence of the Iran Neolithic genetic component (Fernandes et al. 2020), although not to the extent of the Central Italian Imperial period, and the subsequent IA autochthonous people of the Sicani seems to be in genetic continuity with Bronze Age populations (Supplementary figure 1) (Reitsema et al. 2022). It is, however, among the few samples available from the Greek colony of Himera that it is possible to observe individuals with a genetic make-up extremely similar to Villa Falgari and the Imperial individuals (Supplementary table 6) (Reitsema et al. 2022). These individuals may represent the ones that, with their migration from Southern Italy, contributed to formation of the late Republican and Imperial gene pool. It is therefore possible that *Magna Graecia* did not influence only the cultural aspects of the Roman world, but also it greatly contributed to its population dynamics. With the extensive analysis of new Bronze and Iron Age individuals from Southern Italy, especially from a Greek cultural background, it will be possible to clearly indicate the origin of the Near Eastern ancestry in Central Italy eventually discriminating between the different hypotheses.

Italy has always been a place of great genetic variability in the European context (Raveane et al. 2019; Aneli et al. 2021), and the Iron Age/Republican period does not represent an exception (Antonio et al. 2019; Posth et al. 2021; Aneli et al. 2022; Moots et al. 2023; Ravasini et al. 2024). However, it is clear that the variability is also on a temporal dimension rather than only geographical. In Central Italy, the gene pool varies from being more similar to Central Europe, in the first centuries of the Iron Age, to a higher diversity shifting towards the Near East in the late Republican phase, possibly resulting more similar to what it has always been in Southern Italy. For this genetically heterogeneous region, therefore, thinking in broad chronological terms (e.g., Iron Age, Bronze Age etc.) may be simplistic, not allowing for a thorough study of all its complexity. For this particular case given the results of this study, Villa Falgari and the other coeval Near Eastern ancestry individuals should be considered an integrating part of the late Republican gene pool rather than “genetic outliers”, since they may represent the genetic origin of the subsequent periods.

## Methods summary

DNA extraction and library preparation were performed in a dedicated aDNA facility at the University of Tartu, Estonia, and sequencing was performed with Illumina technology following established protocols. After sequencing, data were processed including the steps of trimming, mapping to the human reference genome GRCh37 (hs37d5), check for aDNA damage (Jónsson et al. 2013), contamination estimates (Rasmussen et al. 2011; Korneliussen et al. 2014; Jones et al. 2015), sexing (Skoglund et al. 2013), mitochondrial (Weissensteiner et al. 2016) and Y chromosome (Martiniano et al. 2022) haplogroup assignment, autosomal calling for variants and merging data with the AADR dataset (v54.1 (Dataverse 7.0) Nov 16 2022 https://doi.org/10.7910/DVN/FFIDCW; Mallick et al. 2024) and the data in Reitsema et al. (2022). Subsequently, population genetics analyses were performed including PCA (Patterson et al. 2006), ADMIXTURE analysis (Alexander et al. 2009), D-statistics, outgroup f3-statistics, qpAdm (Patterson et al. 2012; Harney et al. 2021) and DATES analysis (Chintalapati et al. 2022). Detailed methodology is reported in Supplementary Methods.

## Acknowledgements

Bioinformatic analyses were performed with the facilities of the High Performance Computing Center of the University of Tartu. The authors are grateful to the aDNA lab at the Institute of Genomics of the University of Tartu including Lehti Saag, Helja Kabral and Anu Solnik for the support in bioinformatic and wet lab analysis. FR was supported by Sapienza University of Rome grants nr. RM12218167749457 to FC. The EASI-Genomics project has received funding from the European Union’s Horizon 2020 research and innovation programme under grant agreement No 824110.

