## Supplementary Figures for "The arrival of the Near Eastern ancestry in Central Italy predates the onset of the Roman Empire"

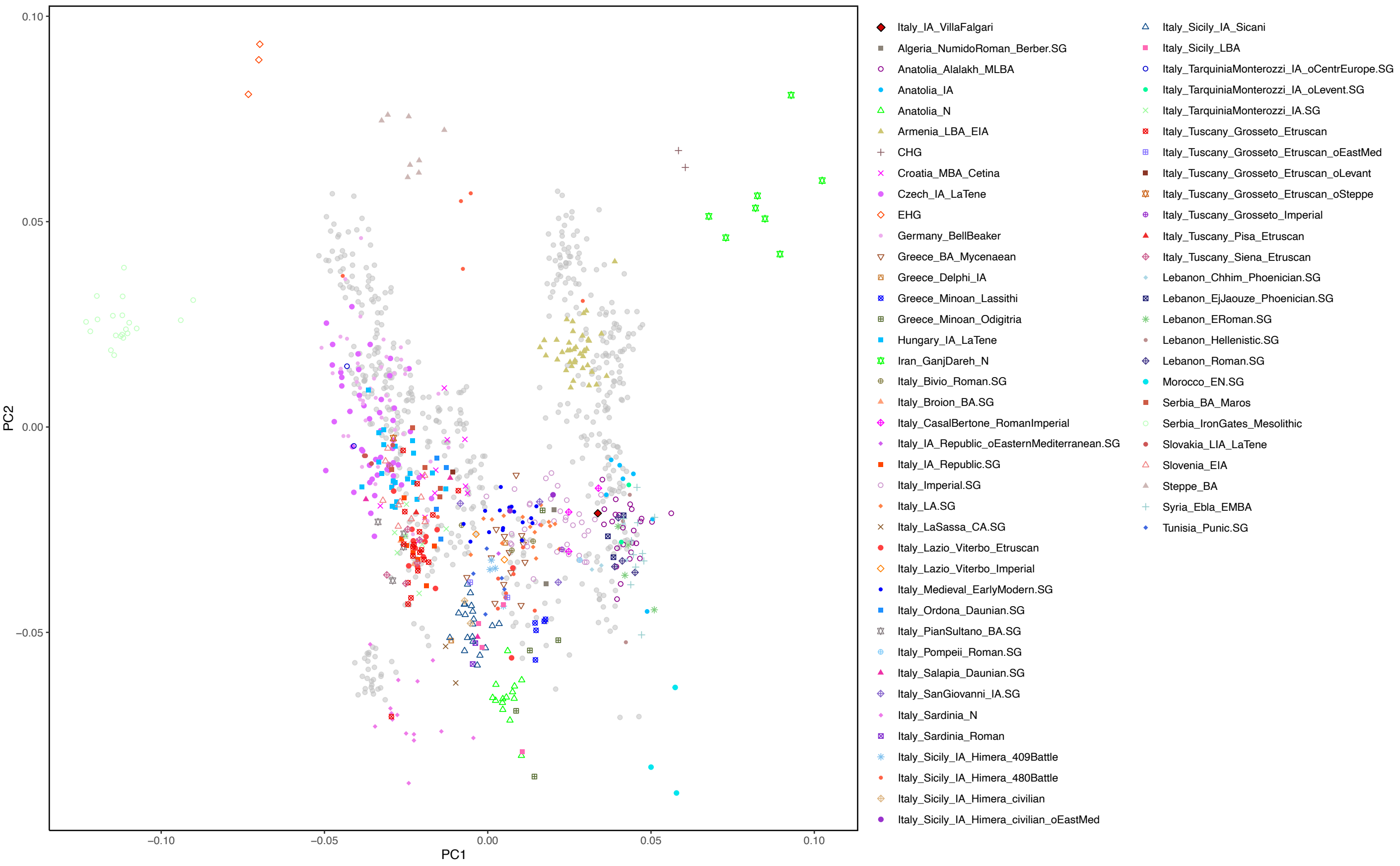


**Supplementary figure 1.** Complete version of the PCA shown in figure 2A. Modern samples are in gray, ancient samples are colored shapes.

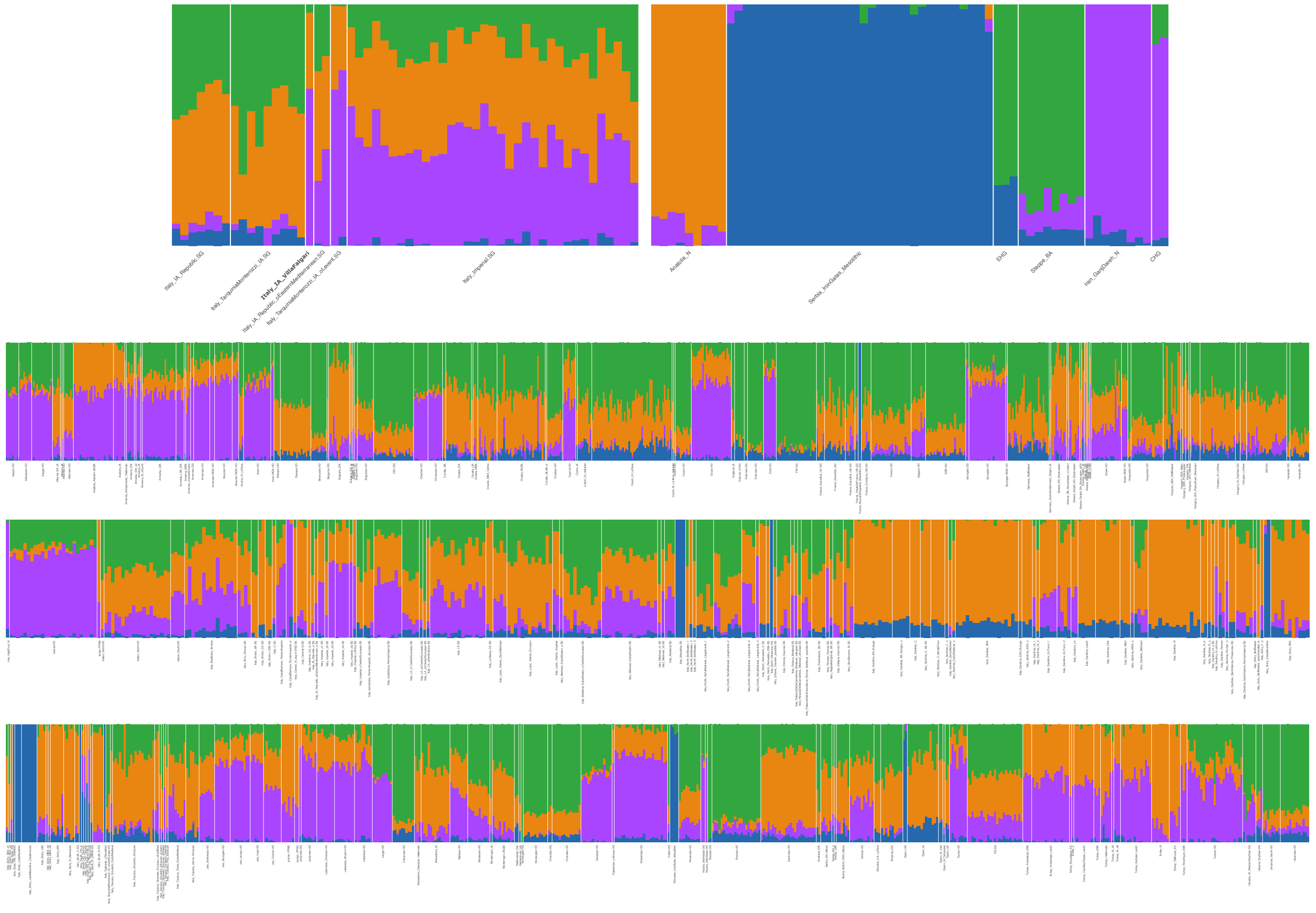


**Supplementary figure 2**. Full version of unsupervised Admixture analysis in figure 2B.

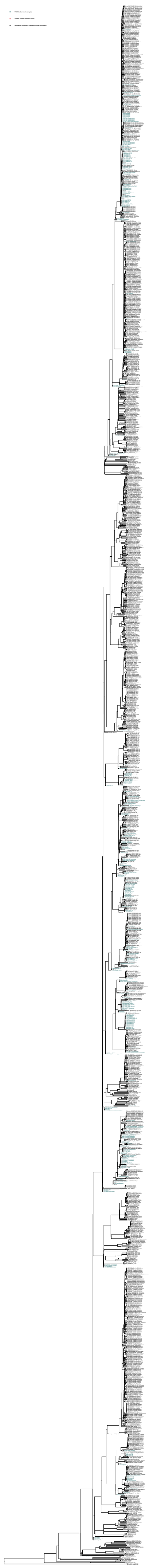

**Supplementary figure 3**. Y chromosome phylogenetic tree reconstructed with pathPhynder. BSP71 is in red, ancient samples from the literature included are in dark green, modern samples in black.

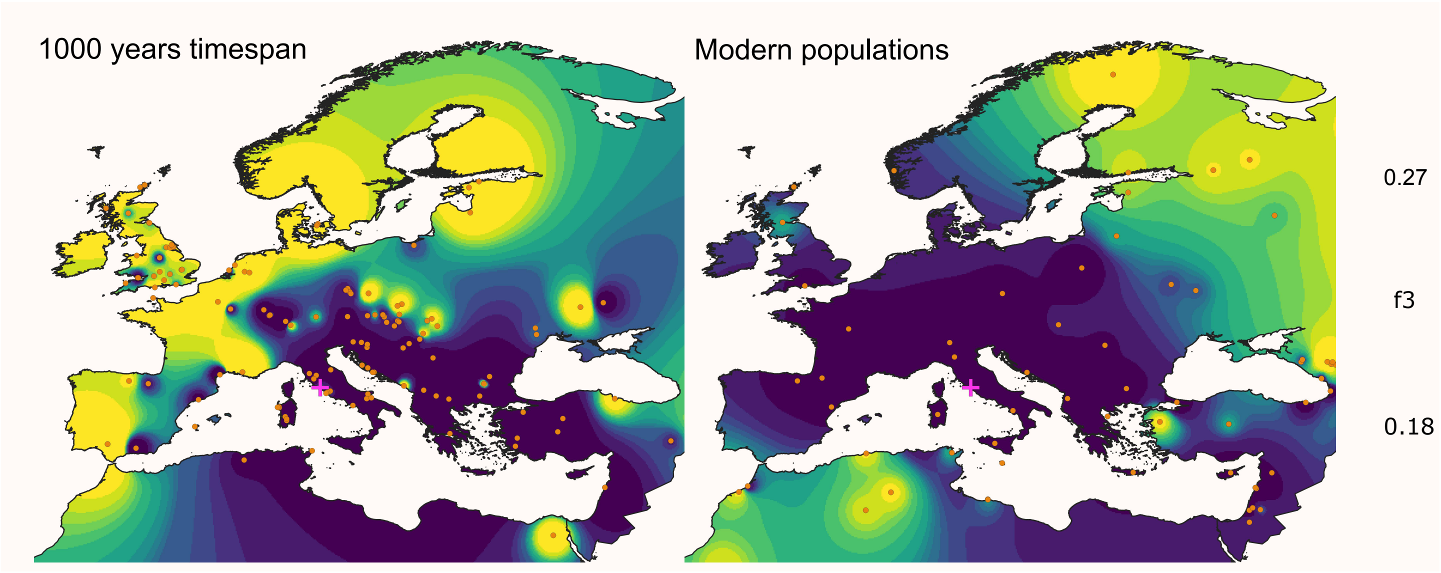


**Supplementary figure 4.** Outgroup f3 statistics interpolated on a European map comparing BSP71 with ancient samples in 1,000 years range from the midpoint of the BSP71 dating (on the left), and with modern populations (on the right).
