## Supplementhary Methods for "The arrival of the Near Eastern ancestry in Central Italy predates the onset of the Roman Empire"

**Supplementary Methods**

Ancient DNA extraction and library preparation were carried out at the aDNA laboratory of the Estonian Biocenter, Institute of Genomics, University of Tartu, Estonia. Quantification and sequencing of the prepared libraries were performed at the Estonian Biocenter Core Laboratory, Institute of Genomics, University of Tartu, Estonia. Bioinformatic analyses were carried out employing the facilities of the High Performance Computing Center of the University of Tartu, Estonia. Radiocarbon dating was performed at the Chrono Centre, Queen's University Belfast, UK.

**Archaeological context**

The individual analyzed in this study comes from Villa Falgari, Tarquinia (Italy) and the archaeological context of this finding is really scarce (Barich et al. 1968). In 1960, an well-preserved almost complete skull was found in this locality without grave goods or any other artifacts allowing for an archaeological dating. Originally, it has been placed in an eneolithic context because of other findings discovered in the same years in the nearby locality of Bandita San Pantaleo (Barich et al. 1968). Nevertheless, recent radiocarbon dating allowed to precisely date this individual to the late Roman Republican phase, suggesting that it might be in some way connected to one of the many other Republican and/or Etruscan necropolises in the area.

**DNA Extraction**

Root portion was taken from a molar tooth of the individual analyzed with a sterile drill wheel, sterilized with 6% (w/v) bleach followed by distilled water and ethanol rinse. The tooth root was placed in 6% (w/v) bleach for 5 minutes, subsequently rinsed with 18.2 MΩcm water for 3 times and soaked for 2 minutes in 70% (v/v) ethanol. The sample was shaken during the bleach and ethanol rinse to allow the detachment of particles. The sample was then positioned on a clean paper towel into a class IIB hood and it was left to dry with the UV light on for two hours. For DNA extraction, the proper volume of EDTA and Proteinase K was calculated based iìon the sample mass, in particular we considered 20x EDTA [µL] of sample mass [mg] and 0.5x Proteinase K [µL] of sample mass [mg]. The sample, EDTA and Proteinase K were placed into a 5 mL PCR-clean conical tube (Eppendorf) under the IIB hood and then incubated on a slow shaker for 72 h at room temperature. DNA extract was concentrated to a final volume of 250 µL with the Vivaspin Turbo 15 (Sartorius). It was later purified in a large volume column using the High Pure Viral Nucleic Acid Large Volume Kit (Roche) with 2.5 mL of PB buffer, 1 mL of PE buffer and 100 µL of EB buffer (MinElute PCR Purification Kit, QIAGEN). The silica column was placed in a collection tube to dry and then in a 1.5 mL DNA lo-bind tube (Eppendorf) for elution. The resulting sample was incubated at 37 °C for 10 minutes with 100 µL of EB buffer and then centrifuged at 13,000 rpm for 2 minutes. The silica column was removed after centrifugation and the sample was stored at -20 °C. 30 µL of the sample was used for library preparation.

**Library preparation and sequencing**

Sequencing library was built with NEBNext DNA library Prep Master Mix Set for 454 (E6070, New England Biolabs) and with Illumina-specific adaptors (Meyer and Kircher 2010) using established protocols (Meyer and Kircher 2010; Orlando et al. 2013; Malaspinas et al. 2014). End repair step was implemented (as described in (Saupe et al. 2021)) using 118.75 µL of water, 7.5 µL of buffer and 3.75 µL of enzyme mix, incubated at 20°C for 30 minutes. The sample was then purified with 500 µL of PB buffer and 650 µL of PE buffer and eluted in 30 µL of EB buffer (MinElute PCR Purification Kit, QIAGEN). Adaptor ligation part was implemented as in Saupe et al. (2021), using 10 µL of buffer, 5 µL of T4 ligase and 5 µL of adaptor mix (Meyer and Kircher 2010), incubating for 14 minutes at 20°C. The sample was purified and then eluted in 30 µL of EB buffer (MinElute PCR Purification Kit, QIAGEN). Adapter fill-in step was carried out using 13 µL of water, 5 µL of buffer and 2 µL of Bst DNA polymerase, incubating for 30 minutes at 37°C and for 20 minutes at 80°C [19]. PCR amplification was performed with 50 µL of DNA library, 1X PCR buffer, 2.5 mM MgCL2, 1 mg/mL BSA, 0.2 µM inPE1.0, 0.2 mM dNTP each, 0.1 U/µL HGS Taq Diamond and 0.2 µM indexing primer. Cycling conditions were settled as follows: 5 s at 94°C, then 18 cycles of 30 s each at 94°C, 60°C and 68°C, and a final extension of 7 minutes at 72°C. Later, the sample was purified with 35 µL of EB buffer (MinElute PCR Purification Kit, QIAGEN). Three verification steps were performed to measure the concentration of dsDNA/sequencing library and to check that library preparation was successful: (1) fluorometric quantification (Qubit, Thermo Fisher Scientific), (2) parallel capillary electrophoresis (Fragment Analyzer, Agilent Technologies) and (3) qPCR. DNA sequencing was performed using the Illumina NextSeq500/550 High-Output single-end 75 cycle kit.

**Mapping**

Before mapping, cutadapt-2.1 (Martin 2011) was employed to remove adapter sequences, indexes, poly-G tails and sequences shorter than 28 bp (--minimum-length 28, to reduce the risk of random mapping of sequences from other species). BWA-0.7.17 (Li and Durbin 2009) was used to map the resulting sequences to the human reference sequence GRCh37 (hs37d5). To mitigate the effect of reference bias in the following analyses we performed mapping with the command bwa aln using relaxed alignment parameters (-n 0.01 -o 2) in combination with disabling seeding (-l 1024) (Kircher 2012; Schubert et al. 2012; Martiniano et al. 2020). The alignment was converted to BAM file format and only the mapped sequences were maintained with samtools-1.9 (Li et al. 2009). Duplicates were removed with picard-2.20.8 (http://broadinstitute.github.io/picard/index.html) and indels were realigned using GATK-3.5 (McKenna et al. 2010). Samtools-1.9 was used to filter out reads with mapping quality lower than 25, as suggested in Martiniano et al. (2020). Reported statistics of the final BAM file are reported in Supplementary Table 1.

**aDNA authentication and contamination rate**

We used the program MapDamage-2.0 (Jónsson et al. 2013) to estimate the frequency 5’ ends of sequences C->T transitions, to check that the sequences obtained are mostly ancient. Contamination rates were estimated on the mitochondrial DNA (mtDNA) with the method detailed in (Jones et al. 2017) by computing the fraction of non-consensus bases at mtDNA haplogroup defining position, and on the X chromosome with the method described in (Rasmussen et al. 2011) implemented in ANGSD (Korneliussen et al. 2014). In both cases contamination estimates were low (<1%) (Supplementary Table 1).

**Genetic sex estimation**

Genetic sex was estimated with the method described in (Skoglund et al. 2013), computing the proportion of reads mapping to the Y chromosome with respect to the total number of reads mapping either to the X or the Y chromosome, setting the R_y limit for males to 0.074.

**mtDNA haplogroups assignment**

Mitochondrial DNA haplogroup was assigned with Haplogrep2 (Weissensteiner et al. 2016). The VCF file used to perform haplogroup assignment was obtained by calling the variants from the BAM files with bcftools-1.14 (Danecek et al. 2021), with the command bcftools mpileup and the additional flag --ignore-RG. Then, only the variant positions were called with the command bcftools call -m --ploidy 1 -v.

**Y chromosome haplogroup assignment and phylogeny**

Y chromosome phylogenetic relationships among the newly reported sample and other ancient samples (Antonio et al. 2019; Lazaridis et al. 2022; Moots et al. 2023) were reconstructed with pathPhynder (Martiniano et al. 2022) starting from BAM files, using standard parameters. As a reference tree, we used the one provided in Martiniano et al. (2022) which spans across all the genetic variability of human Y chromosome haplogroups with a total of 2014 individuals included. To visualize the resulting tree, we used the R packages treeio and ggtree (Yu et al. 2017). The pathPhynder haplogroup assignment has been refined reporting the defining markers for main haplogroups/sub-haplogroups, following the nomenclature proposed by the PhyloTree Y (van Oven et al. 2014) and checking the ISOGG tree v. 15.73.

**Variant calling on autosomes**

Variant calling for autosomes was performed with ANGSD-0.917 (Korneliussen et al. 2014). Haploid genotypes were called sampling a random base (option -doHaploCall 1) for each position present in the 1240K SNPs panel using the -sites options. Major and minor alleles as they are indicated in the 1240K SNPs panel were specified with the option -doMajorMinor 3. The function haploToPlink was used to convert the resulting .haplo file into PLINK (Purcell et al. 2007) format files.

**Design of datasets for genomic analysis**

The Villa Falgari individual was compared to modern and ancient individuals from the literature. Samples included are present in the AADR dataset v54.1 (Dataverse 7.0) Nov 16 2022 <https://doi.org/10.7910/DVN/FFIDCW> (Mallick et al. 2024), both the 1240K and the 1240K+HO dataset, and in (Reitsema et al. 2022). Data coming from different datasets were merged with Plink-1.9. All the population genetics analyses with autosomal data were performed using only transversions to minimize the possible error due to post-mortem damage of ancient DNA.

**Principal component analysis**

For PCA, a dataset of modern and ancient Eurasian and North African samples was selected for a total of 1108 individuals from the literature (Supplementary table 3). PLINK format files were converted into EIGENSTRAT format with convertf of the EIGENSOFT-7.2.0 package (Patterson et al. 2006; Price et al. 2006) using the parameter “familynames:NO”. The program smartpca of the EIGENSOFT-7.2.0 package was used to perform PCA with the parameters lsqproject:YES, autoshrink:YES. Ancient individuals were projected onto the PCs built based on modern samples. The results of the first two PCs were visualized in R-4.1.3 (<https://www.r-project.org/>) with the package ggplot2 (<https://ggplot2.tidyverse.org/>).

**ADMIXTURE analysis**

ADMIXTURE analysis was performed including 1357 modern and ancient Eurasian and North African individuals from the literature (Supplementary table 3). Before running ADMIXTURE, SNPs were pruned for linkage disequilibrium with PLINK (Purcell et al. 2007) (option --indep-pairwise with parameters: 50 5 0.5). Diploid genomes (modern and high-coverage ancient) were converted into pseudo-haploid by selecting a random allele for each variant. 10 independent repetitions (with different random seed, -s option) for k values from 2 to 6 using the option --haploid='*' were performed. The results from different repetitions were merged and visualized with R package pophelper (Francis 2017). k=4 was portrayed because it shows the differentiation between CHG/Iran Neolithic and EHG/Yamnaya components. The name of the ancestral components in the main text (Anatolia Neolithic, Serbia Mesolithic, EHG/Yamnaya, CHG/Iran Neolithic) were given on the basis of the populations considered basal to modern Europeans which have the highest amount of that component.

**Outgroup f3 statistics**

Outgroup f3 statistics was performed in the form f3(Italy_IA_VillaFalgari, Test; ONG.SG). Test populations are relevant modern Eurasians and North Africans or ancient groups in a time interval of 1,000 years, with the mid-date set as the one of Villa Falgari. Longitude and latitude of the Test groups was computed as the mean of the ones of the individuals. f3 statistics were computed with the program qp3pop of the package admixtools-7.0.1 (Patterson et al. 2012) using the option “inbreed:YES”. Interpolation maps with the value of f3 were performed with QGIS-3.26.1 (<https://www.qgis.org/en/site/>). The interpolation was calculated with the IDW method and for representation the option “singleband pseudocolor” was chosen, the minimum value was set to 0.18 and the maximum to 0.27 and, finally, the option “Interpolation: Discrete” was selected.

**D-statistics**

D-statistics was performed in the form D(X, Y; Test, ONG.SG), where X and Y are ancient Italian Iron Age/Republic or Imperial and post-Imperial groups,and Test are ancient Eurasian and North African groups.. D-statistics was performed with the program qpDstat of the package admixtools-7.0.1. (Patterson et al. 2012) with the options “printsd: YES” and “inbreed: YES”.

**qpAdm analysis**

qpAdm analysis was performed with admixtools-7.0.1. (Patterson et al. 2012) with the option “allsnps: YES” in different forms: (1) targets groups were single individuals from Central Italy of the Iron Age/Republican and Imperial period, source populations were the following: Anatolia_N, Iran_GanjDareh_N, Serbia_IronGates_Mesolithic, Steppe_BA; (2) targets were ancient Italian (Iron Age/Republic or Imperial and post-Imperial) groups showing signs of a Near Eastern ancestry, sources were and ancient Italian group showing no major signs of Near Eastern ancestry and a Near Eastern ancient population; (3) targets were post Iron Age/Republican Italian groups, sources were an Iron Age/Republican group with Near Eastern ancestry and a Near Eastern ancient population; (4) targets were post Iron Age/Republican Italian groups, sources were an Iron Age/Republican group with no sign of Near Eastern ancestry and an Iron Age/Republican group with Near Eastern ancestry.

In all four cases the right (outgroup) populations were: Mbuti.DG, Israel_Natufian, Morocco_Iberomaurusian, Iraq_PPNA, Russia_AfontovaGora3, Russia_MA1_HG.SG, Turkey_Boncuklu_N, Turkey_Epipaleolithic, Ethiopia_4500BP.SG, Luxembourg_Loschbour.DG.

**Dates analysis**

DATES (Chintalapati et al. 2022) was employed to direct test admixture dates between a population used as a proxy of the Near Eastern genetic component and the Italian Iron Age gene pool. We used the following parameters: binsize: 0.001; maxdis:1.0; jackknife:YES; qbin:10; runfit:YES; afffit:YES; lovalfit:0.45. We discussed a model including the group Armenia_LBA_EIA as source population for the Near Eastern ancestry because models with sources described in other analysis like PCA and qpAdm (e.g., Iran_N, CHG, Anatolia_IA, Lebanon_Chhim_Phoenician.SG) did not return interpretable results, i.e., negative or extremely (>>200 generations) high admixture dates.
